# Conformable neural interface based on off-stoichiometry thiol-ene-epoxy thermosets

**DOI:** 10.1101/2022.09.22.508978

**Authors:** Eleonora Borda, Marta Jole Ildelfonsa Airaghi Leccardi, Danashi Imani Medagoda, Elodie Geneviève Zollinger, Diego Ghezzi

## Abstract

Off-stoichiometry thiol-ene-epoxy (OSTE+) thermosets have recently gained attention for the rapid prototyping of microfluidic chips because they show low permeability to gases and little absorption of dissolved molecules, they allow direct low-temperature dry bonding without surface treatments, they have a low Young’s modulus, and they can be manufactured via UV polymerisation. The compatibility with standard clean-room processes and the outstanding mechanical properties make OSTE+ an excellent candidate as a novel material for neural implants. Here we exploit OSTE+ to manufacture a conformable multilayer micro-electrocorticography array with 16 platinum electrodes coated with platinum black. The mechanical properties allow device conformability to curved surfaces such as the brain. The low permeability and strong adhesion between layers improve the stability of the device. Acute experiments in mice show the multimodal capacity of the array to record and stimulate the neural tissue by smoothly conforming to the mouse cortex. Devices are not cytotoxic, and immunohistochemistry stainings reveal only modest foreign body reaction after two and six weeks of implantation. This work introduces OSTE+ as a promising material in the field of implantable neural interfaces.

## INTRODUCTION

The nervous system is an extraordinarily complex system capable of receiving, processing and sending a large amount of information simultaneously across multiple central and peripheral regions. Any damage in these areas, caused by illness, genetic disorders or traumatic injuries, might lead to transient or permanent deficits in sensory, cognitive or motor abilities. Advances in neural interfaces [1–6] have contributed to a better understanding of the underlying mechanisms of the nervous system [7,8] and to improving several therapeutic solutions [9–13].

One of the main challenges for neural interfaces is to reduce the foreign body response [14] and preserve their long-term functioning [15,16]. A glial scar is traditionally described as a passive barrier that encapsulates the device, alters the tissue-electrode interface, and impairs neural sensing and stimulation [17–19]. The mechanical mismatch between a neural implant, which is often fabricated with stiff materials, and the soft neural tissue is a crucial limitation inducing neural damage and glial scar formation [20]. The neural tissue is dynamic and subject to deformations caused, for example, by blood pulsation, respiratory pressure, and natural body movements [20,21]. By contrast, most neural interfaces are static. On the other hand, activated microglia and macrophages release soluble factors that can accelerate electrode corrosion and degradation [22]. However, the biological response of the tissue is not the only source of failure. The material and mechanical integrity of the neural interface should also be considered [19]. Electrode corrosion and insulation failure are just two examples that can hinder the long-term stability and performance reliability of these devices [23].

A large body of research in neural interfaces is addressing this issue by investigating novel materials and less invasive surgical approaches. On the contrary, clinical-grade neural implants have undergone only minor changes in the past 70 years [24]. In general, they have large electrode paddles with thick platinum-iridium electrodes and stainless-steel wires embedded in a millimetre-thick silicone matrix [20]. Moreover, a large part of the manufacturing process is done manually. Because of the manufacturing processes and geometrical characteristics, these implants lack important features, such as a high electrode density to improve spatial resolution [25], and low implant stiffness to match the elastic modulus of the neural tissue [26]. These limitations have motivated researchers and newly-founded neurotechnology ventures to focus on flexible thin polymer-based neural interfaces, which conform to curved surfaces and are mechanically compliant with the neural tissue [27–33]. Polyimide (PI), parylene-C and SU-8 are widely used in neural implants because they are biochemically and thermally stable, and processable with silicon-based microfabrication technologies [34–36]. Ultra-thin layers of these materials improved device flexibility achieving relatively long-term integration within the neural tissue [26]. However, when excessively bent, they permanently fold, preventing a tight interface with the nervous tissue [37]. Moreover, the relatively high Young’s modulus (GPa range) and limited elastic deformation [38] make them less suitable for complying with the small and large movements of the nervous tissue [39]. Elastomers such as polydimethylsiloxane (PDMS), on the other hand, exhibit a Young’s modulus three orders of magnitude lower (MPa range) and a much broader elastic regime under strain than PI, parylene-C and SU-8. PDMS has been widely used for conformable and soft interfaces placed on the surface of the spinal cord, brain, retina, and peripheral nerves [38,40–44]. However, despite the superior mechanical properties, micropatterning on PDMS with conventional clean-room processes is an open challenge [45]. Most recently, ultra-soft materials (kPa range), like hydrogels, have also been investigated not only as a conductive coating for electrodes [46] but also as biocompatible substrates matching the mechanical properties of the neural tissue [47,48]. Similarly to PDMS, the microstructuring process of hydrogels and their integration with functional materials are still at an early stage [49]. Moreover, it is still unclear if reducing Young’s modulus of neural implants to this degree would have any benefits in chronically implanted devices [50].

So far, materials for neural implants have been characterised either by excellent compliance with clean-room processes (e.g. PI, parylene-C and SU-8) but high mechanical mismatch with the nervous tissue or, on the contrary, high mechanical compliance but poor manufacturing compatibility (e.g. PDMS and hydrogels). Off-stoichiometry thiol-ene-epoxy (OSTE+) thermosets have been developed for the rapid prototyping of biocompatible microfluidic chips with low permeability to gases and little absorption of dissolved molecules [51–53]. OSTE+ is based on the versatile UV-curable thiol–ene chemistry which provides off-stoichiometry ratios to enable one-step surface modifications, tunable mechanical properties (from GPa to MPa at physiological temperature), and leakage-free sealing via direct UV-bonding [54,55]. In this way, OSTE+ films are patterned with photolithographic techniques and covalently linked via the epoxy- and thiol-binding chemistry without any adhesion enhancing method (e.g. plasma activation). The versatile properties of OSTE+ make them a promising material for neural interfaces [54] encompassing the features of previously used materials, such as excellent compliance with clean-room processes (e.g. PI, parylene-C and SU-8), high mechanical compliance (e.g. PDMS) and, in addition, low permeability to gases and molecules.

Based on the aforementioned reasons, we exploited OSTE+ to fabricate a conformable multilayer micro-electrocorticography (µECoG) array. We characterised mechanical and electrochemical properties as well as showed the multimodal capacity of the implant to record and stimulate the brain in-vivo. Then, we performed an immunohistochemical analysis after two and six weeks of implantation to assess the foreign body response caused by the implanted conformable multilayer OSTE+ µECoG array. This work shows that OSTE+ is a promising material for implantable neural interfaces.

## MATERIALS AND METHODS

### OSTE+

OSTEMER 324 Flex (Mercene Labs) was prepared by mixing the two components in a 1.24:1 ratio.

### Mechanical characterization

OSTEMER 324 Flex mix was spin-coated (1000 rpm, 60 s) onto 4-inch silicon (Si) wafers previously coated with a sacrificial layer in poly(4-styrene-sulfonic acid) (PSS; 561223, Sigma Aldrich). Thiol-ene photopolymerization was performed under UV light (365 nm, 2 min; Gia-Tec). This step was repeated five times to reach a thickness of 150 µm. Samples were then baked overnight at 95 °C to complete the thiol-epoxy thermal polymerization from the thiol excess. Once fully cured, samples were cut by laser (10 J; WS Turret200, Optec Laser Systems) and released in deionized water. Mechanical properties of OSTEMER 324 Flex samples were determined by dynamic mechanical and thermal analysis (DMTA; DMA Q800, TA Instruments) and tensile testing (MTS Systems Corporation). DMTA was operated using a 150-µm thick sample at a thermal ramping of 0.2 °C s^-1^ and a measurement frequency of 1 Hz. For the tensile test, 150-μm thick dog-bone shaped samples (ASTM D412) were mounted in the MTS grips and the crosshead speed was set at 1% of the length between the grips (in mm s^-1^). The displacement and the corresponding force during the test were recorded automatically using the MTS TestSuite™ TW Software (MTS Systems Corporation). The elastic modulus was then calculated as the slope of the curve between 10% and 30% strain (linear regime) using MATLAB (MathWorks), while the elongation at break was defined as the strain with the highest stress value before fracture. Cyclic stress-strain curves were obtained with the same machine by setting in advance the applied percentage of strain. PI samples for the tensile test were prepared by spin-coating PI (PI2611, HD MicroSystems) on a 4-inch Si wafer (1500 rpm, 60 s), soft-baking at 65 °C (5 min) and 95 °C (5 min), and hard-baking at 200 °C (1 hr) and 300 °C (1 hr) both in a nitrogen atmosphere. Samples were cut by laser and peeled off the wafer. PI thickness was 8 µm.

### Resolution and stability tests

A 30-µm thick layer of OSTEMER 324 Flex was spin-coated (1000 rpm, 60 s) onto a PSS-coated 4-inch Si wafer, cured under UV light (2 min) and baked overnight at 95 °C. Photolithography with a 10-µm thick photoresist (AZ 10XT) was performed to pattern platinum (Pt) lines of different sizes and shapes. Soft-baking was performed at 60 °C to avoid cracks on the photoresist due to the softening of OSTEMER 324 Flex at high temperature. Pt was deposited by DC magnetron sputtering (150 nm) followed by lift-off in propylene glycol monomethyl ether acetate (PGMEA, Sigma-Aldrich) during sonication (20 min). To fully encapsulate the metal layer, OSTEMER 324 Flex was spin-coated (6-µm thick, 2500 rpm, 60 s), cured under UV light (2 min) and baked overnight at 95 °C. Samples were then laser-cut and released in deionized water. PI samples were prepared as follows. A titanium/aluminium (Ti/Al, 10/100 nm) release layer was evaporated directly onto a 4-inch Si wafer. A 12-µm thick PI layer was obtained by spin-coating (1000 rpm, 60 s), soft-baking at 65 °C (5 min) and 95 °C (5 min), and hard-baking at 200 °C (1 hr) and 300 °C (1 hr) both in a nitrogen atmosphere. Ti/Pt lines were fabricated by DC magnetron sputtering (15/300 nm) onto PI treated with oxygen plasma (200 W, 20 s), followed by photolithography and chlorine-based dry etching (Corial 210 RIE). The wafer was coated again with a 6-μm thick layer of PI (2000 rpm, 60 s). Samples were cut by laser and released by Al anodic dissolution. Images of the Pt lines were taken with an optical microscope before encapsulation in both OSTEMER 324 Flex and PI samples. The stability test was performed in a sonication bath (185 W) at 37 °C with samples immersed in deionised water.

### Adhesion tests

Samples were designed according to the standard test method for peel resistance and adhesives (T-Peel Test, ASTM D1876-08). Structures were 77.5-mm long and 8.3-mm wide. Two samples were manufactured: one to test adhesion between two OSTEMER 324 Flex layers and the other to test between OSTEMER 324 Flex and Pt. In the first sample, two layers of OSTEMER 324 Flex were prepared as previously described (layer thickness of 150 µm). In the second sample, Pt was deposited by DC magnetron sputtering (150 nm) followed by lift-off in PGMEA during sonication (20 min). In both samples, a 25-µm thick PI foil (Kapton) was used to separate the two layers of OSTEMER 324 Flex. Mechanical T-peel tests were performed using the tensile-load frame with a 10 N load cell at a crosshead translation rate of 0.1 mm s^-1^. The displacement and the corresponding force during the test were recorded automatically using the MTS TestSuite™ TW Software. Tests were performed at room temperature.

### µECoG array fabrication

Multilayer OSTE+ µECoG arrays were designed to cover a large surface of the mouse cortex. A PSS sacrificial layer was deposited on a 4-inch Si wafer, followed by spin-coating of a 30-µm thick layer of OSTEMER 324 Flex as previously described. Photolithography was performed to pattern Pt as previously described. Electrodes (40-µm in diameter) and 30-µm wide feedlines with horseshoe shape (θ = 45 °, W = 30 µm, R = 90 µm) were manufactured on the bottom layer. A 6-μm thick layer of OSTEMER 324 Flex was spin-coated for encapsulation onto the wafer treated with a short and low power oxygen plasma (30 W, 30 s) and exposed to UV laser (375 nm, 800 mJ cm^-2^) with a maskless aligner (MLA 150, Heidelberg). The OSTEMER 324 Flex layer was developed in ethyl L-lactate (77367, Sigma-Aldrich) for 210 s, rinsed in isopropanol and deionised water, dried with a nitrogen gun, and cured at 95 °C overnight. The metallization and OSTEMER 324 Flex encapsulation steps were repeated for the top layer. Multilayer OSTE+ µECoG arrays were then shaped by laser and released from the wafer by PSS dissolution in deionised water. After release, the multilayer OSTE+ µECoG arrays were inserted into a zero insertion force (ZIF) connector placed on a customised printed circuit board. Last, the electrodes were electroplated with platinum black (Pt-black), using a solution containing 1% of platinum chloride (H_2_PtCl_6_ · 6H_2_O), 0.01% of lead acetate (Pb(COOCH_3_)_2_ · 3H_2_O) and 0.0025% of hydrochloric acid (HCl). An LCR meter (4263A, Hewlett Packard) was used for deposition at 800 mV and 100 Hz [56].

### Bending test

Samples for the bending test were fabricated following the same process described for the multilayer OSTE+ µECoG arrays. A customised cycling stretcher was used to perform the bending test with a cyclic compressive force (bending radius: 9.1 mm) [57]. The system included a stepping motor and dedicated mechanics to bend the array. For each test, the stepping motor frequency was set at 0.5 Hz, which resulted in a cycling speed of 10 mm s^-1^. The resistance of the lines was acquired at each cycle and relevant points were then extracted.

### Electrochemical characterization

Electrochemical impedance spectroscopy (EIS) was performed on multilayer OSTE+ µECoG arrays with a potentiostat (CompactStat, Ivium Technologies). The arrays were soaked in phosphate-buffered saline (PBS, pH 7.4) at room temperature with a Pt wire (counter electrode) and an Ag/AgCl wire (reference electrode). The potential was set at 50 mV, and the impedance magnitude and phase were measured between 1 Hz and 100 kHz. Cyclic voltammetry (CV) was performed with the same setup used for EIS. The applied voltage was scanned between −0.6 and 0.8 V at a rate of 50 mV s^−1^ for six cycles. The first cycle was discarded, and the remaining five were averaged.

### Animal handling

All experiments were conducted according to the animal autorisation GE/193/19 approved by the Département de l’emploi, des affaires sociales et de la santé (DEAS), Direction générale de la santé de la République et Canton de Genève in Switzerland. All the experiments were carried out during the day cycle. For the entire duration of the experiment, the health condition was evaluated three times a week, and the body weight was controlled once a week. Experiments were performed in adult (> 1-month-old) female C57BL/6J mice (Charles River Laboratories). Mice were kept in a 12 hr day/night cycle with access to food and water ad libitum. White light (300 ± 50 lux) was present from 7 AM to 7 PM and red light (650-720 nm, 80-100 lux) from 7 PM to 7 AM.

### Acute surgery

Mice were anaesthetized with an intraperitoneal injection of ketamine (87.5 mg kg^-1^) and xylazine (12.5 mg kg^-1^) mixture. Analgesia was performed by subcutaneous injection of buprenorphine (0.1 mg kg^-1^) and lidocaine (6 mg kg^-1^). Artificial tears were used to prevent the eyes from drying. The temperature was maintained at 37 °C with a heating pad. The depth of anaesthesia was assessed with the pedal reflex. The skin of the head was shaved and cleaned with betadine. Mice were then placed on a stereotaxic frame, and the skin was opened to expose the skull. A squared craniotomy was performed to expose the cortex from bregma to lambda landmarks covering both hemispheres.

### Acute recording of epileptiform activity

Immediately after surgery, the multilayer OSTE+ µECoG array was placed on the cortex and the animal was removed from the stereotaxic apparatus while still under anaesthesia. The multilayer OSTE+ µECoG array was then connected to the amplifier using a 32-channel analogue head-stage (Intan Technologies). Signals were recorded using the Open Ephys acquisition board and graphical user interface with the sampling frequency set at 30 kHz [58]. After a baseline period (5 min), 20 µl of pentylenetetrazol (PTZ; 50 mg ml^-1^, 45 mg kg^-1^) were injected intraperitoneally to evoke epileptiform activity [59]. Welch’s power spectral density (PSD) between 0 and 100 Hz was computed in MATLAB.

### Acute brain stimulation and electromyography

Immediately after surgery, the multilayer OSTE+ µECoG array was placed on the cortex to elicit muscle contraction assessed via electromyography (EMG). Two intramuscular needles (working and reference electrodes) were placed in the gastrocnemius muscle with a ground needle in the contralateral back area. Two electrodes of the implant, covering the contralateral cortical hindlimb motor area, were used to deliver bipolar cathodic-first biphasic current pulses. Different pulse durations and amplitudes were tested with ten trials per condition at a repetition rate of 1 Hz. EMG signals were amplified (BM623, Biomedica Mangoni), filtered (1-1000 Hz with 50-Hz notch) and digitalised (16384 Hz). Data were analysed in MATLAB. Data was detrended and bandpass filtered (10 and 1000 Hz). To obtain the EMG envelope, the full-wave was rectified then smoothed using the root-mean-square over a 20-ms window. The integral of the EMG envelope (iEMG) was computed to quantify the total muscle activity.

### Chronic implantation

Mice were anaesthetised with an intraperitoneal injection of ketamine (87.5 mg kg^-1^) and xylazine (12.5 mg kg^-1^) mixture. Analgesia was performed by subcutaneous injection of buprenorphine (0.1 mg kg^-1^) and lidocaine (6 mg kg^-1^). Artificial tears were used to prevent the eyes from drying. The temperature was maintained at 37 °C with a heating pad. The depth of anaesthesia was assessed with the pedal reflex. The skin of the head was shaved and cleaned with betadine. Mice were then placed on a stereotaxic frame, and the skin was opened to expose the skull. A squared craniotomy was performed to expose the left cortex from bregma to lambda. After craniotomy, animals were divided into sham and implanted groups. The implanted group received the multilayer OSTE+ µECoG array below the dura. A sterile cap was placed in contact with the skull bone and glued to it from the outside using bone cement. The sterile cap was filled with a silicone polymer (Kwik-Sil, World Precision Instruments). The skin was then sutured around the cap. The sham group did not receive the implant but followed the same surgical protocol, including the sterile cap.

### Histology

Two and six weeks after implantation, animals were euthanized with an injection of pentobarbital (150 mg kg^-1^). The chest cavity was opened to expose the beating heart, and a needle was inserted in the left ventricle, while the right atrium was cut to allow complete bleeding. The animal was immediately perfused with PBS followed by a fixative solution of 4% paraformaldehyde (PFA) in PBS. At the end of the procedure, the brain was collected and placed in 4% PFA overnight for post-fixation. Brains were cryoprotected in sucrose 30% and frozen in the optimal cutting temperature compound. 30-μm thick coronal sections of the brain were obtained using a cryostat (Histocom) and placed on microscope slides. Brain sections were washed in PBS, permeabilized with PBS + Triton 0.1% (Sigma-Aldrich), left for 1 hr at room temperature in blocking buffer (PBS + Triton 0.1% + normal goat serum 5%), and incubated overnight at 4 °C with primary antibodies for the glial fibrillary acidic protein (GFAP; 1:1000, Z0334, Dako), ionised calcium-binding adapter molecule 1 (IBA1; 1:500, 019-19741, Wako) and neural nuclei (NeuN; 1:500, ABN90P, Sigma). The day after, the sections were incubated for 2 hr at room temperature with secondary antibodies (Alexa Fluor 647, 1:500), counterstained with DAPI (1:300, Sigma-Aldrich) and mounted for imaging with Fluoromount solution (Sigma-Aldrich). Images were acquired at 20x with a slide scanner microscope (VS120, Olympus). Image quantification was performed in ImageJ. First, a rectangular region of interest was manually selected to represent the area in contact with the implant, and the corresponding area on the other hemisphere. Each image was converted to binary using a threshold algorithm: all pixels whose intensity was above the threshold were assigned the value 1, while the value 0 was assigned to the rest of the pixels. For the three markers, the threshold was defined using the Moment thresholding method in ImageJ. The percentage of the pixels above the threshold was then computed.

### Cytotoxicity

A test on extract was performed on multilayer OSTE+ µECoG arrays sterilised by UV exposure. Arrays were incubated in water for 0 (no incubation), 1, 2, 5, 6, and 8 days before the test. For each condition, two arrays were used and the test on each array was replicated three times. The extraction on the samples was performed for 24 hr at 37 °C and 5% CO_2_ with 1 ml of extraction vehicle for 3 cm^2^ of device surface. The extraction vehicle was Eagle’s minimum essential medium (11090081, Thermo Fisher Scientific) supplemented with 10% foetal bovine serum (10270106, Thermo Fisher Scientific), 1% penicillin-streptomycin (15070063, Thermo Fisher Scientific), 2 mM L-Glutamine (25030081, Thermo Fisher Scientific) and 2.50 μg ml^-1^ Amphotericin B (15290026, Thermo Fisher Scientific). L929 cells (88102702, Sigma-Aldrich) were plated in a 96-well plate at a subconfluent density of 7000 cells per well in 100 μl of the same medium. L929 cells were incubated for 24 hr at 37 °C and 5% CO_2_. After incubation, the medium was removed from the cells and replaced with the extract (100 μl per well). After another incubation of 24 hr, 50 μl per well of XTT reagent (Cell proliferation kit 11465015001, Sigma-Aldrich) were added and incubated for 4 hr at 37 °C and 5% CO_2_. An aliquot of 100 μl was then transferred from each well into the corresponding well of a new plate, and the optical density was measured at 450 nm by using a plate reader (FlexStation3, MolecularDevices). Clean medium alone was used as a negative control, whereas medium supplemented with 15% of dimethyl sulfoxide (D2650-5×5ML, Sigma-Aldrich) was used as a positive control. Optical density was normalised to the average value of the negative control to determine the cell viability.

### Statistical analysis and graphical representation

Statistical analysis and graphical representation were performed with Prism 8 (Graph Pad). The D’Agostino & Pearson omnibus normality test was performed to justify the choice of the statistical test. In plots, p-values were reported as: * p < 0.05, ** p < 0.01, *** p < 0.001, and **** p < 0.0001.

## RESULTS AND DISCUSSION

### OSTE+ manufacturing

Among the commercially available OSTE+ polymers, we chose OSTEMER 324 Flex, which differs from other OSTE+ resins in its mechanical properties that make it conformable. After mixing the three monomers with thiol, allyl, and epoxy groups, the viscous resin can be coated onto a wafer surface and the thickness of the layer can be controlled by the spin speed (**Figure 1a**). Then, the fast thiol-ene radical UV polymerization results in a soft solid surface, which is chemically reactive due to the exposed thiol excess and epoxy groups. Last, the slower anionic thermal polymerization allows epoxy groups to react with the remaining thiol groups, thereby stiffening the substrate [54,60,61]. UV polymerisation allows photo-patterning of the film, which acts as a negative photoresist (**Figure 1b**). OSTE+ allows precise patterning of various sizes and shapes with relatively high resolution (about 2 µm). The possibility of UV patterning of OSTE+ is an advantage for the encapsulation step of neural implants if compared to PDMS encapsulation, which is usually done manually [38,40,62] or by reactive-ion etching to obtain smaller features [42] but with the risk of leaving toxic residues [63].

**Figure 1.**
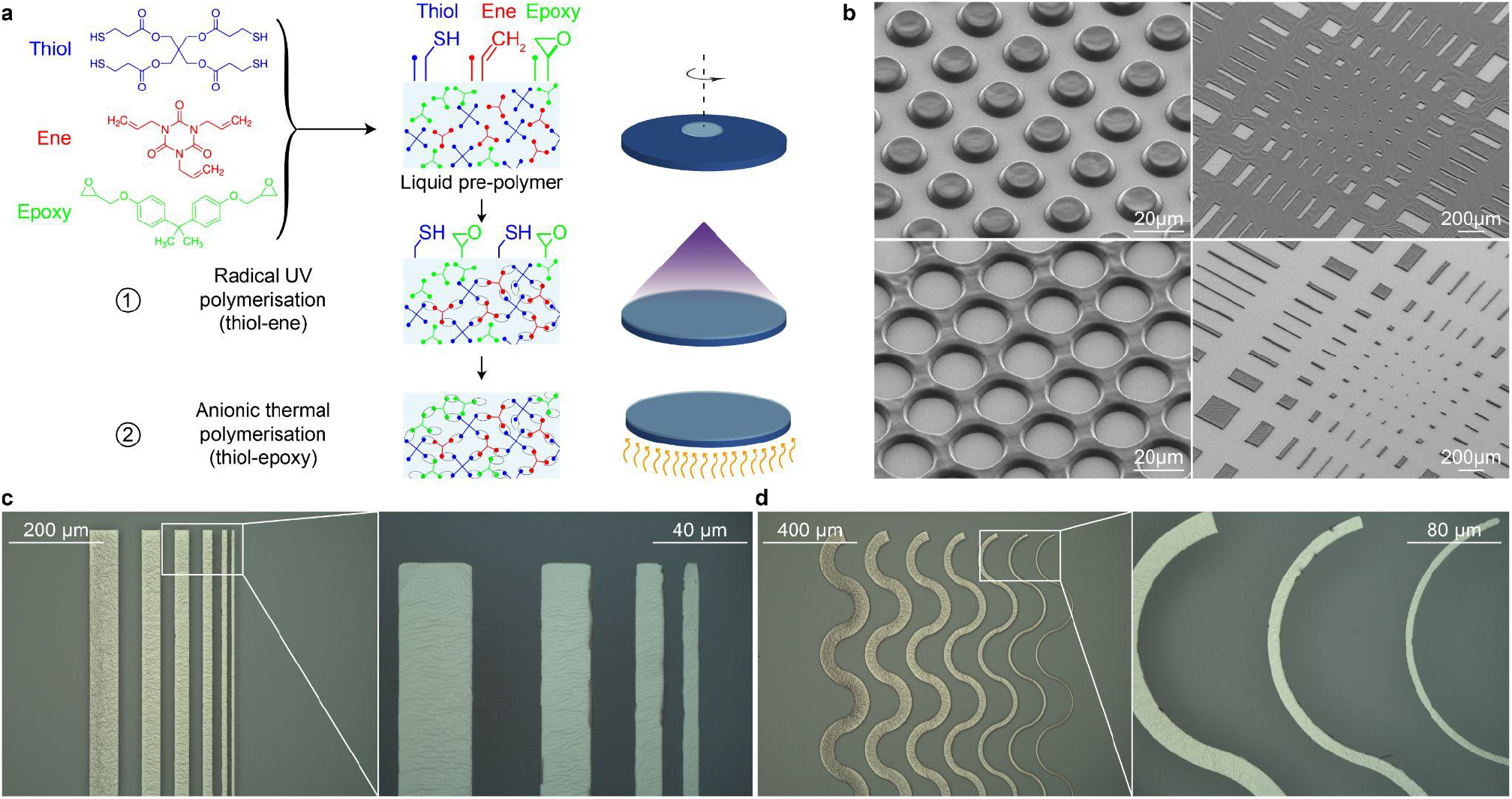
OSTE+ manufacturing. (**a**) Step-by-step procedure to manufacture a spin-coated film of OSTE+. (**b**) Example of structures obtained with OSTEMER 324 Flex via direct UV exposure of the liquid pre-polymer. The layer thickness is 4 µm. (**c,d**) Resolution of straight (**c**) and horseshoe-shaped (**d**) lines on OSTEMER 324 Flex. Line widths are 60 µm, 40 µm, 30 µm, 20 µm, 10 µm and 5 µm from left to right. The magnifications show line widths 30 µm to 5 µm (**c**) and 20 µm to 5 µm (**d**).

Next, we assessed the resolution of thin metal films on OSTEMER 324 Flex obtained with standard photolithographic techniques. Straight (**Figure 1c**) and horseshoe-shaped (**Figure 1d**) lines of decreasing width from 60 µm down to 5 µm have been patterned with 10-µm thick photoresist. 5-µm lines appears sharply defined on the OSTEMER 324 Flex substrate, which is a largely better resolution that what typically obtained on PDMS [42,64]. Ripples appeared on the Pt layer because of the stress that builds up between two materials (Pt and OSTEMER 324 Flex) of different modulus and coefficients of thermal expansion, like happening on other elastomeric substrates including PDMS [65,66]. Ripples help increase elastic deformation in the neural interface. Similarly to PI, parylene-C and SU-8, the possibility of using standard photolithography to directly pattern metallic structures ensures high reliability and reproducibility in the fabrication process. Unlike previous studies using thiol-ene-based polymers for neural interfaces, the process presented here is simple and does not require any transfer-by-polymerization process [63,67–70].

### OSTE+ mechanical characterization

A crucial element for materials used in neural interfaces is flexibility and conformability. Ultra-thin layers of PI, parylene-C and SU-8 allows device flexibility. However, the relatively high Young’s modulus (GPa range) and limited elastic deformation make them less attractive than elastomers like PDMS (MPa range). OSTEMER 324 Flex closes this gap in neural interfaces having promising mechanical properties.

First, we performed DMTA in one OSTEMER 324 Flex sample (**Figure 2a**). The glass transition temperature (T_g_) is 29.2 °C (**Figure 2a**, green circle), lower than the operating body temperature for neural implants (37 °C). This feature is interesting compared to other materials (PI, parylene-C, SU-8 and PDMS) because OSTEMER 324 Flex is stiffer at room temperature (easier insertion) and softens once inserted into the body (reduced mechanical mismatch) [67,68,71,72]. Accordingly, the storage moduli at room (21 °C) and physiological (37 °C) temperatures are respectively 323.7 MPa and 15.2 MPa (**Figure 2a**, red circles). Then, we performed tensile stress/strain tests (**Figure 2b**) to quantify the elastic modulus (**Figure 2c**), stress at break (**Figure 2d**), and elongation at break (**Figure 2e**). OSTEMER 324 Flex samples have an average Elastic modulus of 15.03 ± 1.21 MPa, an average stress at break of 15.70 ± 4.95 MPa and an average elongation at break of 65.88 ± 10.37 % (n = 4 samples, mean ± s.d.).

**Figure 2.**
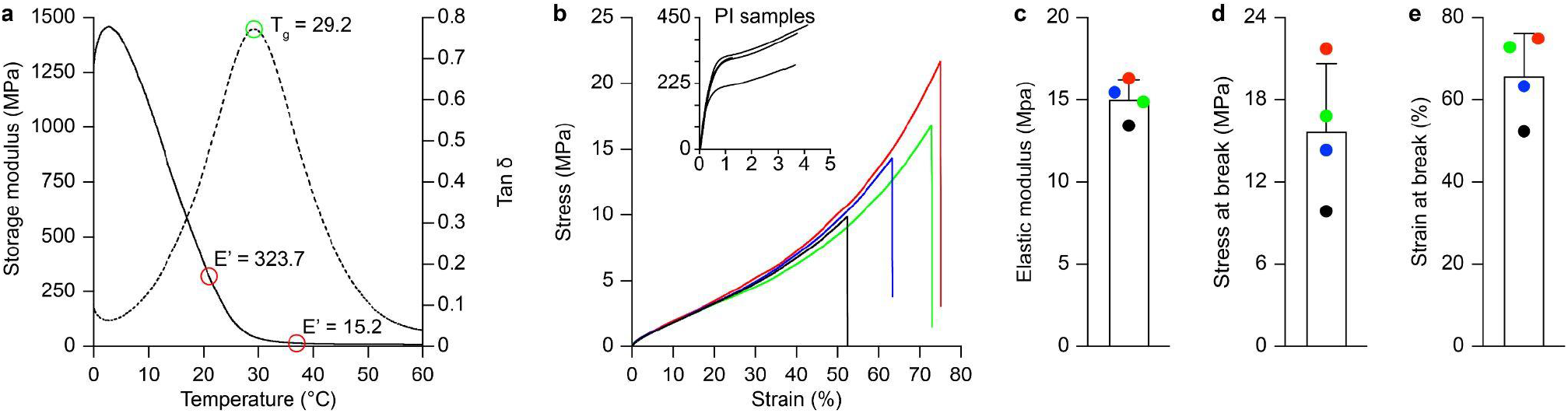
OSTE+ mechanical characterization. (**a**) DMTA in one OSTEMER 324 Flex sample. The black line is the storage modulus and the dashed line is the tan δ as a function of the temperature. The green circle highlights the T_g_, and the two red circles highlight the storage modulus at 21 and 37 °C. (**b**) Stress-strain curves of four OSTEMER 324 Flex samples under tensile test at room temperature. The insert shows the stress-strain curves obtained from four PI samples. (**c-e**) Quantification of the elastic modulus (**c**), stress at break (**d**) and strain at break (**e**) from the four tested samples (n = 4 samples, mean ± s.d.).

For µECoG arrays, it is important to take into consideration the capability of a flexible polymeric substrate to conform with the complex topography of the central nervous system. Therefore, we computed the critical thickness (h) to achieve spontaneous wrapping of an OSTEMER 324 Flex film around a curvilinear body with a given radius of curvature, following equation (1) for an elasto-capillary model [35,73].

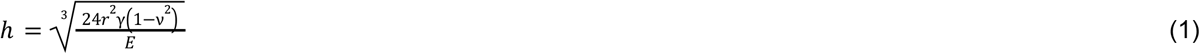

The radius of curvature r is 2.3 mm, corresponding to the mouse cortex [35], the elastic modulus E is 15.03 × 10^6^ N m^−2^, the Poisson’s ratio v is assumed 0.4 [74], and the cerebrospinal fluid surface tension γ is 61 mN m^−1^ [75]. The threshold value for conformability to a mouse brain is 75.64 µm. Below this thickness, multilayer OSTE+ µECoG arrays naturally conform to the mouse brain without additional pressure. In addition, because of its elastic component, OSTEMER 324 Flex, can conform better than PI to curved surfaces like spheres. For comparison, results from tensile stress/strain tests in conformable PI samples (8-µm thick) are reported in the insert of **Figure 2b**.

Based on these results, OSTEMER 324 Flex is a suitable candidate for surface and penetrating brain implants, where the dynamic strain is in the range of 0.01-0.03% [76], or intraneural interfaces for nerves away from joints [77].

### OSTE+ adhesion strength

The adhesion between layers is a crucial feature to ensure integrity and longevity of a neural interface. A common failure mechanism in neural implants is the delamination between the insulating layers and the conductor, leading to fluid penetration and loss of the device integrity and functionality [78]. Therefore, we performed adhesion T-peel tests to investigate the adhesion strength of OSTEMER 324 Flex samples (**Figure 3**). First, we assessed the self-adhesion forces between two OSTEMER 324 Flex layers (**Figure 3a**). During a tensile load, the peel force linearly increases with the distance until cohesive fracture occurs in the unbonded portion of the sample before delamination. This result suggests a strong self-adhesion because of the OSTE+ functional groups covalently linked during the anionic thermal polymerisation [55]. Moreover, heating the polymer above the T_g_ during the anionic thermal curing softens the polymer and allows it conforming to nano and microcavities in the substrate creating a perfect seal. In a neural interface, it is also crucial to verify the adhesion strength also between the encapsulation and the metal electrodes. Therefore, we repeated the T-peel test in samples with a Pt layer between the two OSTEMER 324 Flex layers (**Figure 3b**). In this case, the top OSTE+ layer was peeled off because of the weaker adhesion to Pt at an averaged peel force per unit width of 89.22 ± 8.11 N m^-1^ (mean ± s.d., n = 4 samples), which is 30 times higher than the one required to peel Ti sputtered on PI [78]. The peel force per unit width during separation was averaged between 1 and 5 mm of displacement. Ti is commonly used as an adhesion layer for Pt in PI-based implants; OSTEMER 324 Flex showed stronger adhesion to Pt without the need of any intermediated interface nor surface treatment.

**Figure 3.**
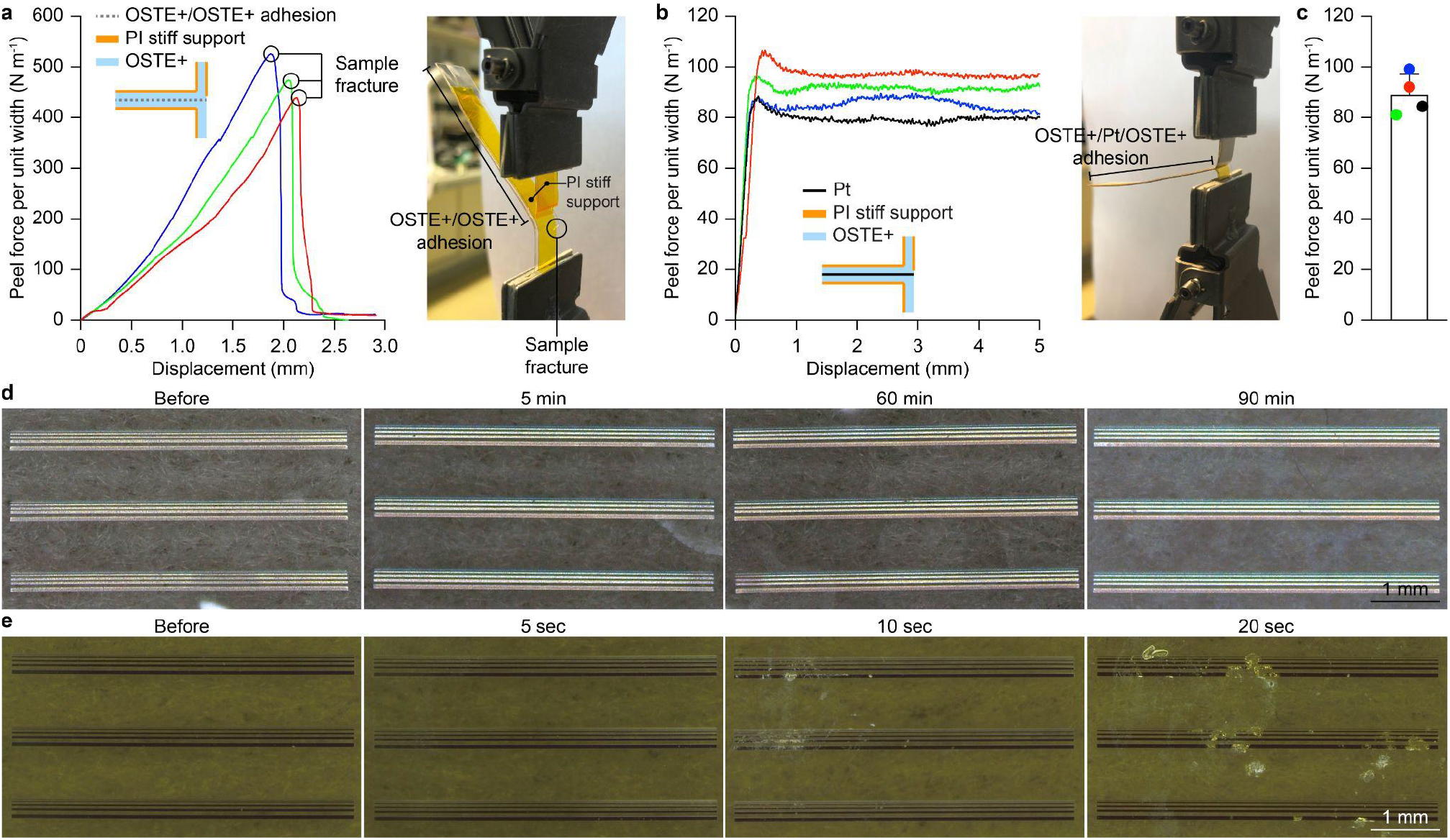
OSTE+ adhesion strength. (**a**) T-peel test between two OSTEMER 324 Flex layers. In all three samples a fracture of the samples is observed. The insert represents the cross-section of the sample. The image on the right shows a sample under testing. (**b**) T-peel test between two OSTEMER 324 Flex layers with a Pt film in between (4 samples). The insert represents the cross-section of the sample. The image on the right shows a sample under testing. (**c**) Quantification of the averaged peel force per unit width at which layers separate. Each colour corresponds to the samples in (**b**). (**d**) Stability test for OSTEMER 324 Flex. Images before sonication and after 5, 60 and 90 min of sonication. (**e**) Stability test for PI. Images before sonication and after 5, 10 and 20 s of sonication.

Then, we fabricated samples with patterned Pt lines fully embedded in either two layers of OSTEMER 324 Flex or PI to compare the encapsulation capability of the materials. Samples were then immersed in a sonication bath and controlled at various time points. The PI sample shows signs of metal delamination already after 10 s of sonication (**Figure 3e**) while the OSTE+ samples remain unaltered even after 90 min (**Figure 3d**). This qualitative test demonstrates the superior stability of OSTEMER 324 Flex.

### OSTE+ electrical and electrochemical characterization

Next, we designed and optimised a process-flow to fabricate conformable multilayer OSTE+ µECoG arrays by using established wafer-scale technologies. The implant is designed to covers a large area of the mouse cortex (**Figure 4a,b**) and 16 electrodes are arranged to cover both brain hemispheres (eight electrodes per side) placed in two layers: half in the bottom layer (**Figure 4a**, black) and half in the top layer (**Figure 4a**, grey). Multilayer fabrication is advantageous to not increase the area occupied by feedlines or to manufacture three-dimensional electrodes [56,79]. The total thickness of the device is 42 µm, well below the critical thickness for conformability. At this thickness, OSTEMER 324 Flex has a bending stiffness (B) of 1.10 × 10^−7^ N m, according to equation (2) [35].

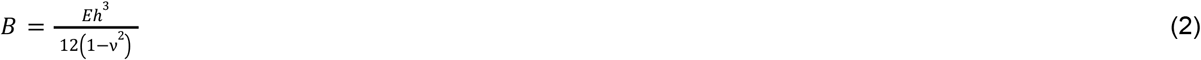

This value is lower than the bending stiffness for 8 µm-thick PI (4.34 × 10^−7^ N m) and 150 µm-thick PDMS (3.75 × 10^−7^ N m), while a 42-µm thick OSTE+ array is still easy to handle.

**Figure 4.**
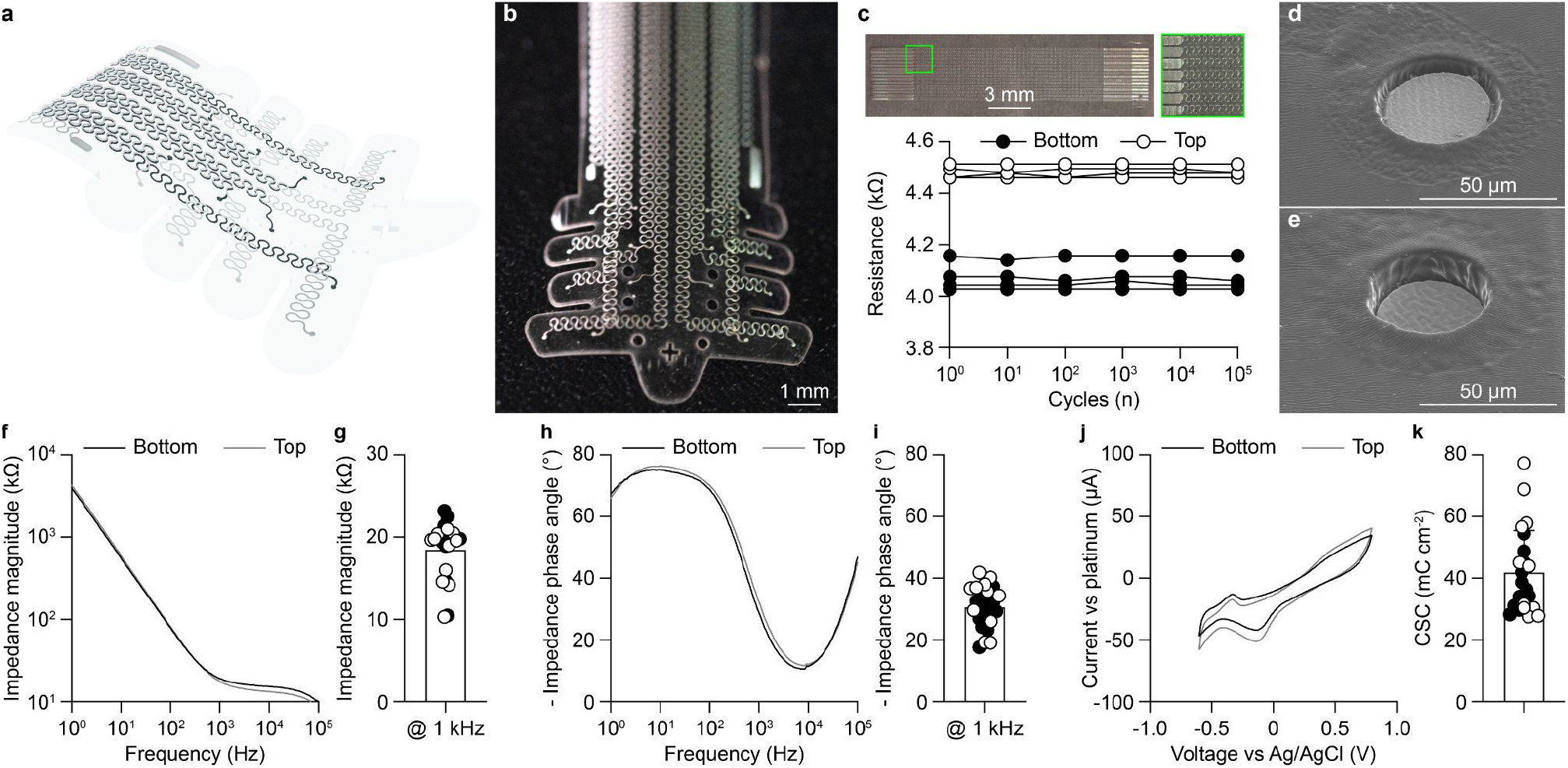
OSTE+ electrical and electrochemical characterisation. (**a**) Sketch of conformable multilayer OSTE+ µECoG array highlighting the bottom (black) and top (grey) electrode layers. (**b**) Picture of the conformable multilayer OSTE+ µECoG array. (**c**) Line resistance as a function of multiple bending cycles (bending radius: 9.1 mm). The white dots are top electrodes (n = 4), whereas the black dots are bottom electrodes (n = 4). The top pictures show the sample used for the bending test. (**d,e**) Scanning electron microscopy images of 40-µm diameter top (**d**) and bottom (**e**) electrodes coated with Pt-black and encapsulated in OSTE+. (**f**) Plot of the impedance magnitude. The gray line is the average from the top electrodes (n = 11 electrodes from N = 2 arrays) and the black line is the average from the bottom electrodes (n = 12 from N = 2 arrays). (**g**) Quantification (mean ± s.d.) of the impedance magnitude at 1 kHz. The black circles are bottom electrodes and the white cirlces are top electrodes (p = 0.3237, two-tailed Mann-Whitney test). (**h**) Plot of the impedance phase angle. The gray line is the average from the top electrodes (n = 11 electrodes from N = 2 arrays) and the black line is the average from the bottom electrodes (n = 12 from N = 2 arrays). (**i**) Quantification (mean ± s.d.) of the impedance phase angle at 1 kHz. The black circles are bottom electrodes and the white cirlces are top electrodes (p = 0.2239, two-tailed unpaired t test). (**j**) Plot of the CV. The gray line is the average from the top electrodes (n = 11 electrodes from N = 2 arrays) and the black line is the average from the bottom electrodes (n = 12 from N = 2 arrays). (**k**) Quantification (mean ± s.d.) of the CSC. The black circles are bottom electrodes and the white cirlces are top electrodes (p = 0.2738, two-tailed unpaired t test).

The substrate is 30 µm and each encapsulation is 6 µm. Therefore, metal lines are not located in the neutral plane, potentially causing stress concentration upon bending and torsion during handling and implantation [80]. Hence, a horseshoe pattern was used to reduce the stress in the stiff metal layer as well as allow conformability to the tissue (**Figure 4a,b**). Meander-like patterns have already been investigated by other groups, who found the horseshoe pattern superior over sinusoidal or U-shaped designs [80–82]. We performed measurements of the electrical resistance of the horseshoe lines under bending cycles to evaluate their robustness (**Figure 4c**). The resistance values of bottom and top lines remained stable at least for 10^5^ bending cycles. However, it can be noticed a consistent but small difference between the two layers. The top layer has a 10.05% increase compared to the bottom layer. This small difference might be caused by changes in the Pt microstructure happening on the bottom layer due to the additional heat treatments following the fabrication of the top layer [62], or by small changes in the deposition chamber between the two processes. If, on the one hand, a horseshoe pattern provides flexibility, on the other hand, it reduces the integration density and slightly increases the electrical resistivity. Alternatively, stretchable conductors could be used [38,42].

Direct writing photolithography was used to pattern the electrode openings in both layers (**Figure 4d,e**). The UV polymerization of OSTE+ allows multilayer fabrication and reduces the risk of misalignment, thus ensuring a stable and repeatable process. Electrodes have been coated with Pt-black, which is known to reduce their impedance magnitude as well as increase the total charge storage capacity (CSC) compared to bare Pt [56]. Moreover, no significant difference has been found in the electrochemical properties between bottom and top electrodes (**Figure 4f-k**). The versatility of the fabrication process allows electrode coating with other materials, like sputtered iridium oxide, spin-coated or electrodeposited PEDOT:PSS or a combination of the two [35].

### OSTE+ functional validation in-vivo

Conformable multilayer OSTE+ µECoG arrays were tested for brain recording and stimulation.

First, we performed in-vivo brain recordings in an anaesthetised mouse upon induction of epileptiform activity. The conformable multilayer OSTE+ µECoG array was implanted with the electrodes covering both hemispheres of the cortex from lambda to bregma (**Figure 5a,d**). The neural activity was recorded before and after peritoneal injection of the convulsant PTZ, which is routinely used to test anticonvulsants in animals [59,83–85]. The transition between baseline (**Figure 5a**, black) and epileptiform activity (**Figure 5a**, red) after PTZ injection was clearly detected by the conformable multilayer OSTE+ µECoG array with an excellent signal to noise ratio, thus demonstrating the potential of OSTE+ neural interfaces for monitoring brain activity. Injection of PTZ induced characteristic spike and wave discharges [86], evident in all the 16 spectrograms from top and bottom electrodes (**Figure 5b)**. The presence of epileptiform activity is further evidenced by the statistically higher PSD that was observed after PTZ injection (**Figure 5c**).

**Figure 5.**
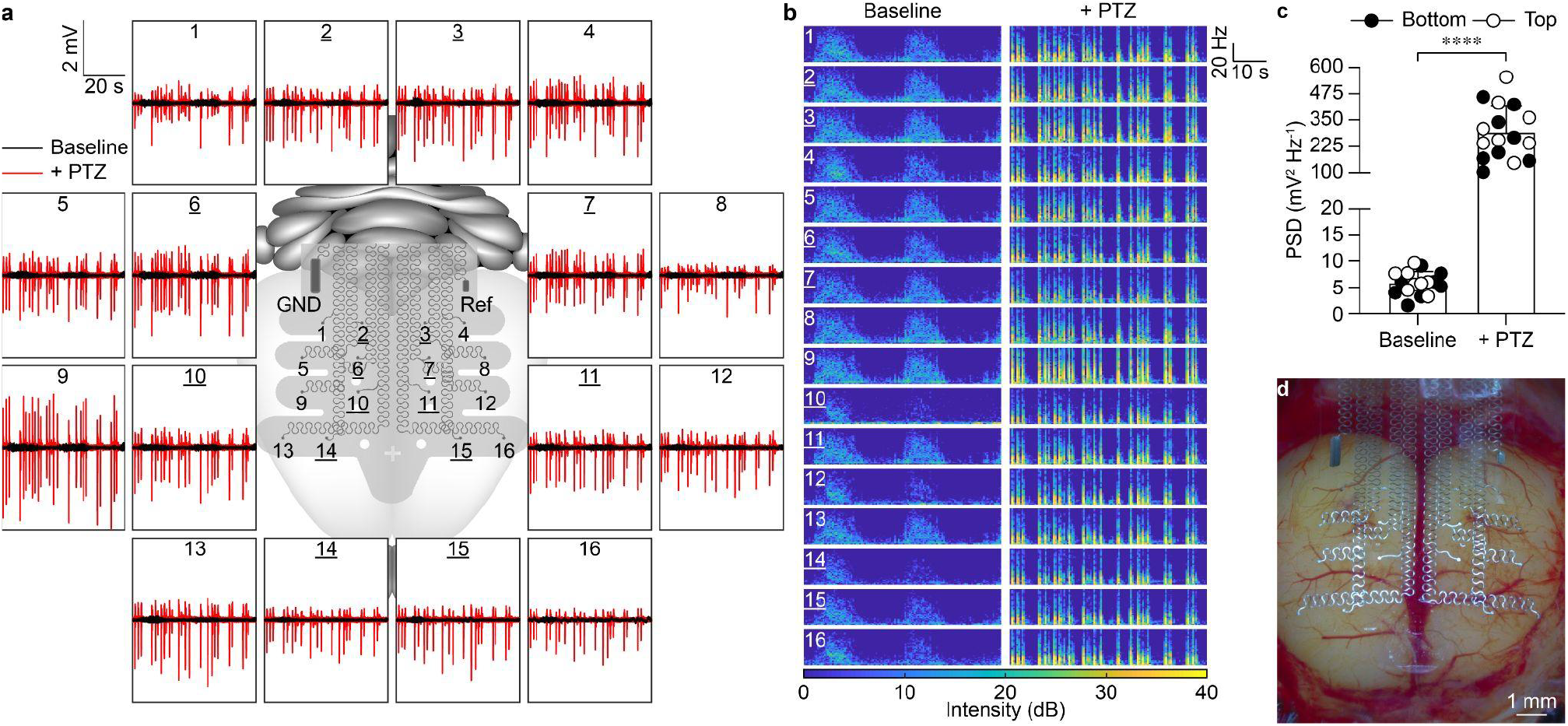
Monitoring of epileptiform activity. (**a**) The central sketch shows a multilayer OSTE+ ECoG array covering a mouse brain for in-vivo recordings. Medial electrodes (underlined numbers) are in the bottom layer. Around the sketch, representative recordings from each electrode are reported. The black traces are the baseline activity, while the red traces are the epileptiform activity after PTZ injection. Each box corresponds to 60 seconds of recordings. (**b**) Spectrogram between 0 and 50 Hz from the 16 recordings shown in **a**. (**c**) PSD quantification between 0 and 100 Hz from recordings in **a**. The black circles are bottom electrodes and the white circles are top electrodes (p < 0.0001, two-tailed paired t test). (**d**) Image of the multilayer OSTE+ ECoG array onto the mouse cortex.

Next, we investigated the performance of conformable multilayer OSTE+ µECoG arrays in brain stimulation in one anaesthetised mouse (**Figure 6**). Two neighbouring electrodes covering the cortical hindlimb motor area have been used for bipolar electrical stimulation to elicit muscle contraction assessed via EMG recordings (**Figure 6a**). Two intramuscular needles were placed in the gastrocnemius muscle, while a ground needle in the contralateral back area. EMG responses appeared upon single cathodic-first biphasic pulses (**Figure 6b**). Then, we extracted the linear envelope of the EMG signal (**Figure 6c**) and computed the iEMG to compare various current amplitudes and pulse durations (**Figure 6d**). As expected, longer pulse durations reduced the current required for inducing robust motor activity.

**Figure 6.**
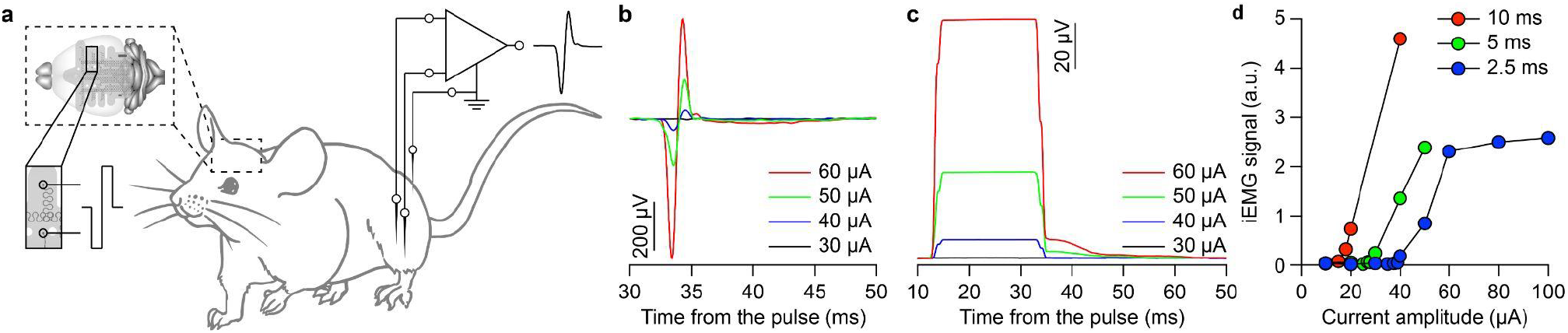
Stimulation of the cortical hindlimb motor area. (**a**) Diagram of the stimulation set-up. A pair of electrodes in the µECoG array covering the cortical hindlimb motor area are selected for stimulation. Two needles are in the mouse hindlimb for EMG recordings. A third needle is the ground. (**b**) EMG responses to increasing current amplitudes from 30 to 60 uA with a 2.5-ms long pulse. (**c**) EMG envelope calculated using the root-mean square over a 20-ms sliding window from responses in **b**. (**d**) Quantification of the iEMG for 2.5 (blue), 5 (green) and 10 (red) ms pulses at increasing current amplitudes.

Overall, we showed that conformable multilayer OSTE+ µECoG arrays are suitable for acute brain recordings and stimulation of neural activity. In future works, in-vivo long-term functionality of conformable multilayer OSTE+ µECoG arrays will necessarily be assessed.

### OSTE+ biocompatibility

Last, we evaluated the OSTE+ biocompatibility in-vitro and in-vivo.

Previous studies showed that OSTE+ films are non-toxic to cultured cells only after immersion in water for 7 days [61]. Therefore, we assessed the in-vitro cytotoxicity of conformable multilayer OSTE+ µECoG arrays after immersion in water for 1, 2, 5, 6, and 8 days before the test or without immersion (0 days). The results showed no cytotoxicity for all the tested conditions, including the arrays not immersed in water before the test (Figure 7**a,b**). This result might be explained by the several washing steps performed during fabrication which could remove impurities from the residual constituents.

**Figure 7.**
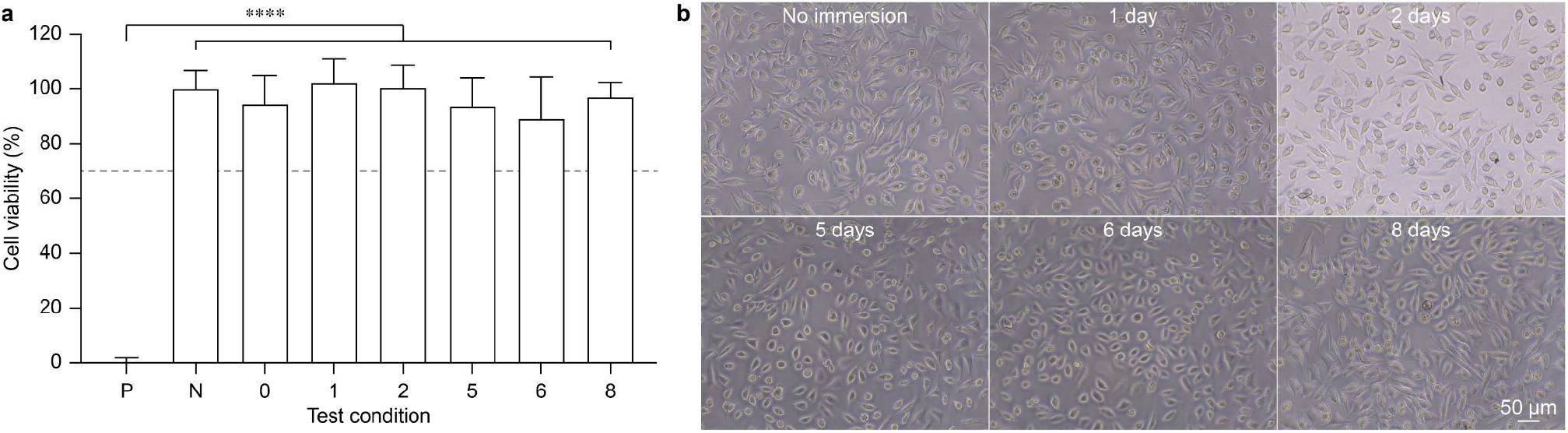
OSTE+ in-vitro cytotoxicity. (**a**) Quantification of cell viability in the conditions tested (mean ± s.d.). Positive control (P) 0 ± 1.8% (9 replicas); negative control (N) 100 ± 6.8% (9 replicas); no immersion (0) 94.5 ± 10.5% (2 arrays, 6 replicas); 1 day immersion (1) 102.2 ± 8.8% (2 arrays, 6 replicas); 2 days immersion (2) 100.4 ± 8.4% (2 arrays, 6 replicas); 5 days immersion (5) 93.7 ± 10.5% (2 arrays, 6 replicas); 6 days immersion (6) 89.0 ± 15.6% (2 arrays, 6 replicas); 8 days immersion (8) 97.0 ± 5.3% (2 arrays, 6 replicas). One-way ANOVA: F = 144.1 and p < 0.0001. Tukey’s multiple comparisons test: P vs. all other conditions p < 0.0001; all other comparisons are statistically not significant (p = 0.3516 or higher). The dashed grey bar indicates the threshold for cytoxicity. (**b**) Representative optical images of the cells after the test, for different immersion conditions.

Then, we evaluated the in-vivo biocompatibility of conformable multilayer OSTE+ µECoG arrays by performing chronic implantation and post-mortem histological assays against GFAP, IBA1 and NeuN markers (**Figure 8**). For this analysis, conformable multilayer OSTE+ µECoG arrays have been implanted in one brain hemisphere (operated), leaving the other hemisphere untouched (unoperated). Histological analysis has been performed on coronal brain sections corresponding to the operated area at two time points: two and six weeks after implantation. Images at both two and six weeks post implantation were quantified and statistically compared with a Two-Way repeated measure ANOVA (Factor 1 is the procedure: implanted or sham; Factor 2 is the brain hemisphere: operated or unoperated). For NeuN signals, there are no statistically significant differences at both two (Factor 1: p = 0.4329; Factor 2: p = 0.5359; Interaction: p = 0.3888) and six (Factor 1: p = 0.2382; Factor 2: p = 0.7065; Interaction: p = 0.4628) weeks post implantation (**Figure 8a,b**). Similarly, IBA1 signals are not significantly different at both two (Factor 1: p = 0.4331; Factor 2: p = 0.0701; Interaction: p = 0.1705) and six (Factor 1: p = 0.5754; Factor 2: p = 0.5902; Interaction: p = 0.8532) weeks post implantation (**Figure 8a,d**). GFAP signals are not significantly different at two weeks post implantation (Factor 1: p = 0.4438; Factor 2: p = 0.1805; Interaction: p = 0.4333), but they show a statistically significant difference after six weeks (Factor 1: p = 0.0044; Factor 2: p = 0.2612; Interaction: p = 0.4430) between implanted and sham groups (**Figure 8a,c**). The Šidák’s multiple comparison test showed that this difference is associated to the unoperated hemisphere (unoperated sham vs unpoperated implanted: p = 0.099) and not to the operated one (operated sham vs unoperated implanted: p = 0.1006).

**Figure 8.**
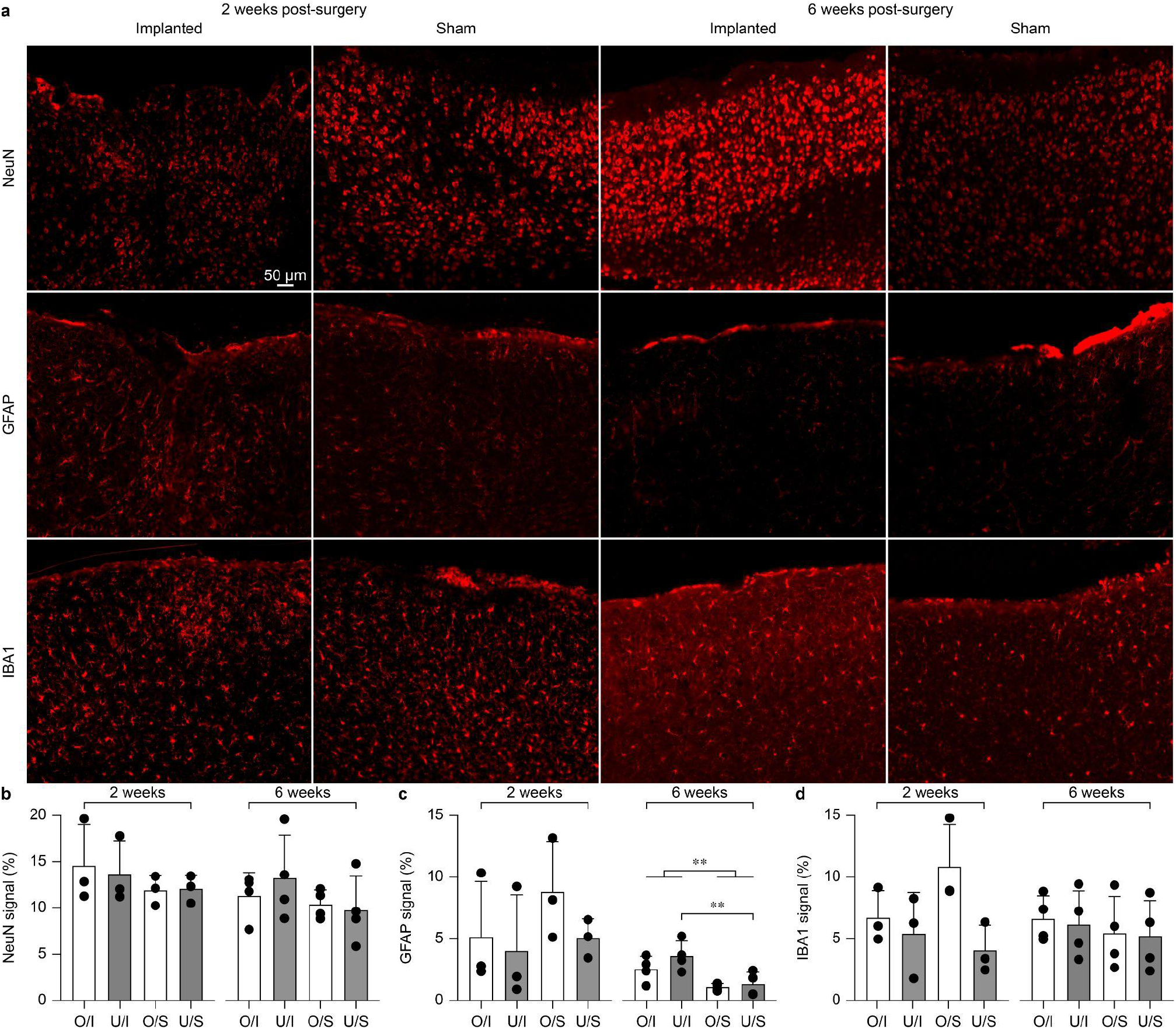
OSTE+ in-vivo histological analysis. (**a**) Representative images for NeuN, GFAP and IBA1 sampled 2 and 6 weeks after implantation in both implanted and sham mice. Corresponding images for NeuN, GFAP and IBA1 are from the same mouse. Slice were sampled at the level of the operated area. (**b-d**) Quantification of the NeuN (**b**), GFAP (**c**) ans IBA1 (**d**) signals two (mean ± s.d., n = 3 mice) and six (mean ± s.d., n = 4 mice) weeks after implantation in implanted and sham mice. O/I: operated hemisphere for implanted mice; U/I: unoperated hemisphere for implanted mice; O/S: operated hemisphere for sham mice; U/S: unoperated hemisphere for sham mice.

From these results, we concluded that superficial damage is caused mainly by the surgical procedure as previously observed [87], and conformable multilayer OSTE+ µECoG arrays do not cause significant foreign body reaction in the first six weeks post implantation.

## CONCLUSIONS

We exploited a soft and flexible OSTE+ formulation, known as OSTEMER 324 Flex, to develop a conformable neural interface. OSTE+ thermosets have shown tunable mechanical properties by adjusting the stoichiometry of the constituting monomers, they can be manufactured via UV polymerisation and allow direct low-temperature dry bonding without surface treatments [55]. These physicochemical properties result in several advantages compared to other materials used in neural interfaces (PI, SU8, parylene-C or PDMS).

OSTE+ shows superior adhesion forces, a necessary feature to improve stability and longevity of neural interfaces [78]. In addition, previous studies have shown that OSTE+ have low permeability to gases and show low absorption of small molecules: two crucial characteristics to protect thin implants against moisture and ions in long-term chronic applications [25].

Conformable multilayer OSTE+ µECoG arrays are manufactured with standard clean-room manufacturing processes. Compared to PDMS, photolithography on OSTE+ allows reliable patterning to make openings for electrical contacts, otherwise done manually [38,40,62] or by aggressive etching steps [42,63]. Achieved resolution is comparable to other polymeric neural interfaces based on PI, parylene-C or SU-8, but OSTEMER 324 Flex showed superior mechanical and encapsulation properties. A glass transition temperature lower than the body temperature might allow temperature-dependent softening during surgical insertion: an interesting feature for penetrating neural implants [67,68,71,72].

The superior adhesive and encapsulation properties of OSTE+ derives from the two-step polymerisation process, which could be further exploited to expand the device functionality. Reactive groups exposed at the surface allow for surface functionalization. For example, to embed molecules improving biocompatibility and reducing long-term foreign body response. Also, the radical UV polymerisation might be exploited to integrate microfluidics in neural implants for drug release [39].

The electrophysiological and histological in-vivo study showed that OSTEMER 324 Flex is a promising material to build conformable multilayer μEcoG arrays. Given these promising results, future works will focus on long-term in-vivo testing.

## ACKNOWLEDGEMENTS

The authors would like to thank the Neural Microsystems Platform at Fondation Campus Biotech Geneva for their support with the optimization of the fabrication process. This work was supported by École Polytechnique Fédérale de Lausanne, Medtronic plc and the Swiss National Science Foundation (200021_182670).

## AUTHOR CONTRIBUTIONS

E.B. fabricated the devices, performed experiments and data analysis, and wrote the manuscript. M.J.I.A.L. designed the fabrication process. D.I.M. performed brain stimulation and data analysis. E.G.Z. performed histological assessment. D.G. led the study and wrote the manuscript. All the authors read and accepted the manuscript.

## DATA AVAILABILITY

The authors declare that all other relevant data supporting the findings of the study are available in this article and in its supplementary information file. Access to our raw data can be obtained from the corresponding author upon reasonable request.

